# Highly localized chemical sampling at sub-second temporal resolution enabled with a silicon nanodialysis platform at exceedingly slow flows

**DOI:** 10.1101/2023.09.08.556607

**Authors:** Insu Park, Sungho Kim, Christopher Kenji Brenden, Weihua Shi, Hrishikesh Iyer, Rashid Bashir, Yurii Vlasov

## Abstract

Microdialysis (MD) is a versatile and powerful technique for chemical profiling of biological tissues and is widely used for quantification of neurotransmitters, neuropeptides, metabolites, biomarkers, and drugs in the central nervous system as well as in dermatology, ophthalmology, and in pain research. However, MD performance is severely limited by fundamental tradeoffs between chemical sensitivity, spatial resolution, and temporal response. Here, by using wafer-scale silicon microfabrication, we develop and demonstrate a nanodialysis (ND) sampling probe that enables highly localized chemical sampling with 100μm spatial resolution and sub-second temporal resolution at high recovery rates. These performance metrics, which are 100X-1000X superior to existing MD approaches, are enabled by a 100X reduction of the microfluidic channel cross-section, a corresponding drastic 100X reduction of flow rates to exceedingly slow few nL/min flows, and integration of a nanometer-thin nanoporous membrane with high transport flux into the probe sampling area. Miniaturized ND probes may allow for the minimally invasive and highly localized sampling and chemical profiling in live biological tissues with unprecedented spatio-temporal resolution for clinical, biomedical, and pharmaceutical applications.

## Introduction

Microdialysis (MD) enables sampling endogenous and exogenous chemicals from the extracellular space (ECS) in biological tissues^1^. MD is based on the diffusion of chemicals from the ECS over a semi-permeable membrane that is continuously flushed with an inlet aqueous flow (perfusate, Fig.1A) to collect analytes in the outlet flow (dialysate, Fig.1A) for subsequent chemical analysis. MD is a widely used method in fundamental and clinical neuroscience and enables quantification of neurotransmitters, neuropeptides, metabolites, biomarkers, and drugs in the central nervous system (CNS)^2^. It is also used widely for monitoring pharmacokinetics and pharmacodynamics in dermatology, ophthalmology, and in pain research^3^. While versatile, the method has known disadvantages related to fundamental tradeoffs between the perfusate flow rate 𝑄, the analytes recovery rate 𝑅, and the probe sampling area 𝑆, which result in a competition between the probe’s chemical sensitivity, spatial resolution, and its temporal resolution^4^. While operation at ultra-slow flow (USF) rates of 1μL/min can produce concentration levels of the analyte in the collected sample approaching that in the ECS (recovery rate 𝑅 approaching 100%), however, typical USF perfusion requires long collection periods to obtain appropriate sample volumes for analytes quantitation. The USF perfusion also significantly decreases the already relatively slow temporal resolution of MD^5^. In addition, in a typical commercial MD probe with a millimeter long membrane wrapped around a capillary of 200−400 μm in diameter, the resulting active membrane surface area 𝑆 is of the order of a few mm^2^, which averages out chemical content over large brain volumes. The typically large probe cross-section (0.15-0.5 mm^2^) also results in inevitable tissue trauma and inflammation during implantation that complicates chemical analysis.

**Figure 1.**
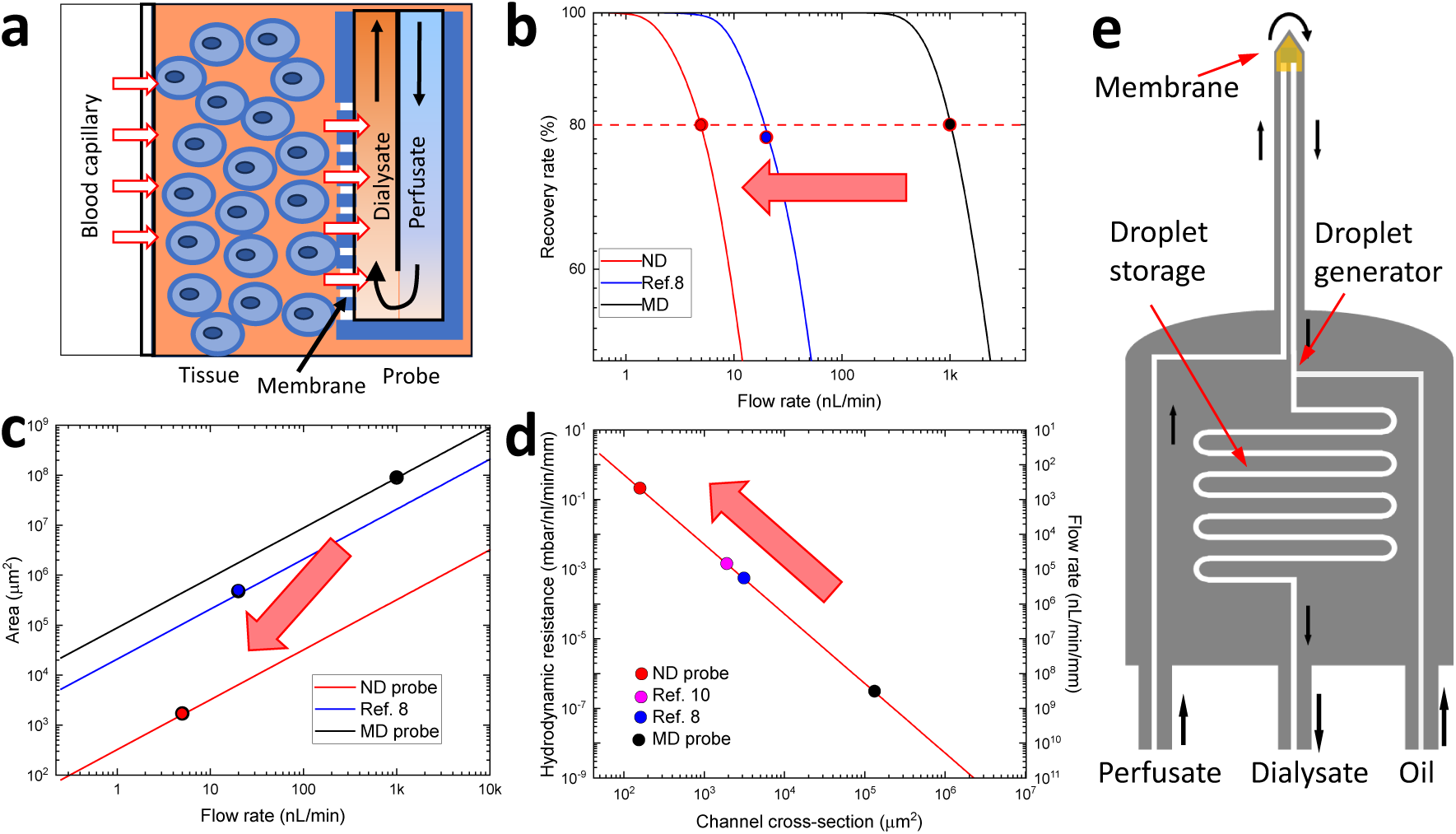
Design tradeoffs for the ultimate ND probe. **A)** Schematics of the dialysis process**. B)** Recovery rate versus perfusate flow rates for various micro- and nano-dialysis probes. Black – commercial MD probe with 𝑆 = 8 · 10^7^ μm^2^ and 𝑅 = 3 · 10^7^ sec/cm, Blue – microfabricated probe of Ref. 8 with 𝑆 = 5 · 10^5^ μm^2^ and 𝑅 = 8 · 10^6^ sec/cm, Red – ND probe with 𝑆 = 1.7 · 10^3^ μm^2^ and 𝑅 = 1 · 10^5^ sec/cm. **C)** Membrane surface area 𝑆 required to achieve 80% recovery rate versus perfusate flow rate for various membrane resistances. Black, blue, and red lines are for 3 · 10^7^ sec/cm, 8 · 10^6^ sec/cm, and 1 · 10^5^ sec/cm, respectively. **D)** Hydrodynamic resistance for water vs microfluidic channel cross-section for various probes. Right vertical axis is perfusate flow rate per 1mm of a channel length for a given differential pressure of 100 mbar **E)** Schematics of an integrated ND probe comprising of an ultrathin small area membrane at the tip of a sampling needle, scaled microfluidic channels, a droplet generator, and a delay line for droplet storage.

Miniaturization of MD probes using microfabrication technologies^6–14^ has begun to address these issues by decreasing the flow rates to 100nL/min and below^8,14^ enabling significant reduction of the sampling area down to just a few thousands of μm^2^ that, together with integration with droplet microfluidics^9–12,14^, results in significantly improved temporal resolution down to a few seconds. What are the limits of this miniaturization? Is it possible to design a microfabricated probe that has a sampling area of just 1000μm^2^, a recovery rate approaching 100%, the capability to manipulate and deliver sample volumes of just 10pL to sensitive chemical analysis methods at 1 second temporal resolution, all the while occupying a cross-section of just 1000μm^2^ to minimize tissue trauma? While some of these numbers, which correspond to a 100X to 10,000X improvement compared to commercial MD probes, have been demonstrated separately, they have never been realized in a single device due to very restrictive fundamental tradeoffs.

Here we show that by utilizing wafer-scale silicon microfabrication a nanodialysis (ND) sampling probe with an integrated ultrathin 30nm nanoporous silicon membrane enables ultra-localized chemical sampling from an area as small as 1700μm^2^. This superior performance is enabled by the drastic reduction of the microfluidic channel cross-section to 40μm^2^ and operation at the exceedingly low flow rates of just a few nL/min. On-chip droplet segmentation of the dialysate flow enables sub-second temporal resolution and an on-chip storage of the segmented analytes in a time-preserved sequential droplet train.

## Results

### Design of the ultimate NanoDialysis (ND) probe

The governing equation which describes the recovery rate 𝑅 of the dialysis probe in the steady-state equilibrium model^4^ is defined as

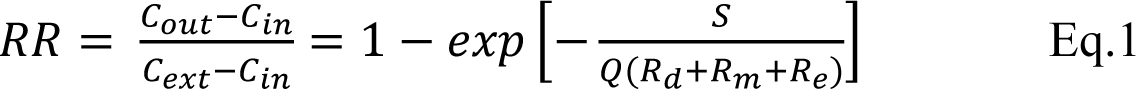

where 𝑆 is the active area of the membrane, 𝑄 is the perfusate flow rate, and 𝑅_𝑑_, 𝑅_𝑚_, 𝑅_𝑒_ are the mass transport resistances for the dialysate, the membrane, and the tissue, respectively. 𝐶_𝑜_, 𝐶_𝑖_, and 𝐶_𝑒𝑒𝑜_ are the concentrations of analyte in the dialysate, in the perfusate (typically zero), and in the undisturbed tissue, respectively.

To achieve the primary design goal of highly localized chemical sampling, the active probe surface area 𝑆 has to be minimized while keeping the recovery rate 𝑅 considerably large (e.g., >80%). The corresponding product 𝑆⁄𝑄(𝑅_𝑑_ + 𝑅_𝑚_ + 𝑅_𝑒_) in Eq.1 should be larger than −ln(1 − 0.8) = 1.6 (Fig.1B). Drastic reduction of this product can be achieved by simply removing the membrane from the design and flushing out the surrounding tissue using balanced push and pull flows^7,9–12,14^ (push-pull perfusion, PPP), or just by pulling in the interstitial fluid (ISF) from the ECS with a single open orifice^9^ (ultrafiltration, UF). In all of these implementations, the effective sampling volume is estimated^15,16^ to be still of the order of 3 · 10^6^μm^2^ (or 3nL), which is accompanied by a strong depletion of analytes in the tissue surrounding the probe’s flushing site, hence further complicating the analysis^3^. Furthermore, tissue trauma due to exposure to bulk flow into and out of the probe is inevitable^15^, especially at relatively high μL/min flow rates common to PPP and UF.

An alternative approach to reduce the membrane area by 1000X is to decrease the 𝑅_𝑚_𝑄 product by a corresponding 1000X (Fig.1C) by decreasing the membrane resistance 𝑅_𝑚_ and the flow rate 𝑄. The pore size, porosity, thickness, and tortuosity of a membrane can be adjusted to decrease 𝑅_𝑚_. Polymer-based membranes used in commercial MD probes have relatively high 𝑅_𝑚_ of the order of 𝑅 = 3 · 10^7^ sec/cm due to their large thickness (∼50 μm) and tortuosity, which translates into large areas and high flow rates (Fig.1C, black) to achieve high recovery. A microfabricated MD probe^8^ with an integrated 5μm-thick anodic aluminum oxide (AAO) membrane has been demonstrated, however with just 1% porosity resulting in high 𝑅 of the order of 10^7^ sec/cm and 𝑆 about 10^5^ μm^2^ (Fig.1C, blue). Atomically thin (0.34nm) graphene-based membranes can provide extremely low resistance^17,18^ which is promising for solvent nanofiltration^19^, however their defects-induced pore sizes below 1nm are too small for the commonly targeted molecular weight cut-off (MWCO) of a few kDa. Utilization of molecularly thin (15nm-50nm) nanoporous silicon^20–22^ or silicon nitride membranes^23^ with low resistances can potentially provide high 𝑅 with active surface aresa 𝑆 down to 1000 μm^2^ (Fig.1C, red). This would require, however, reduction of flow rates 𝑄 to the exceedingly slow nL/min range (Fig.1C, red dot) that is at least 100X-1000X slower than flow rates used in traditional USF MD^5^ and in microfabricated probes^8,10,12^.

Such exceedingly slow flow rates impose numerous challenges in flow control, analyte collection time, sample manipulation, and chemical quantitation. These problems can be mitigated, at least in part, by drastic reduction of the microfluidic channel cross-section from 100,000μm^2^ in traditional MD probes to about 2000μm^2^ in the microfabricated MD probes^6,8,9,14^ and even further to channels as small as 50 μm^2^ using silicon microfabrication^24^. Such a dramatic channel down-scaling of channel cross-section results in a corresponding increase of the hydrodynamic resistance (Fig.1D), which is beneficial for maintaining a precise control of nL/min flow rates at reasonable pressures of 100mbar per 1 mm of the channel length.

Hence, to realize the ultimate nanodialysis (ND) probe (Fig.1E) that can be used for highly localized and efficient chemical sampling with minimal tissue damage, the deeply scaled microfluidic channels supporting exceedingly slow flow rates need to be integrated with an ultrasmall sampling area covered with an ultra-thin semipermeable membrane.

### Concept and fabrication of the in-plane open-flow ND probe

Our ultimate goal is to integrate an nm-thin semipermeable membrane with a high density of nanopores into a silicon probe, shown in Fig.1E to achieve a high recovery rate 𝑅 with an ultra-small sampling area 𝑆. Besides improving the 𝑅, decrease of the flow rates to nL/min can help to minimize stress on the already fragile membrane that otherwise would result in its rupture that is often observed at high flow rates^8^ even for membranes as thick as 5μm. To avoid such stress from building up we adopted an in-plane flow design (Fig.2A) where laminar flows into and out of the sampling area are tightly balanced, hence cancelling the out of plane bulk flow. This design is in striking contrast to the more traditional push-pull sampling approaches, where the microfluidic flows are intentionally redirected from the inside of the probe into the tissue^12^ either perpendicular to the in-plane flows inside the probe^7,11,14^ (Fig.2B) or from the capillary located above the sample orifice^9,10^ (Fig.2C). The resulting 𝑅 is increasing, however at the expense of bulk flow inside the tissue, which can cause fluid loss/accumulation as well as tissue damage. Instead, the in-plane flow design minimizes stresses toward the external tissue and/or encapsulating membrane. Our design is therefore, also conceptually different from the open-flow micro-perfusion^3^ and can be called open-flow nanodialysis.

**Figure 2.**
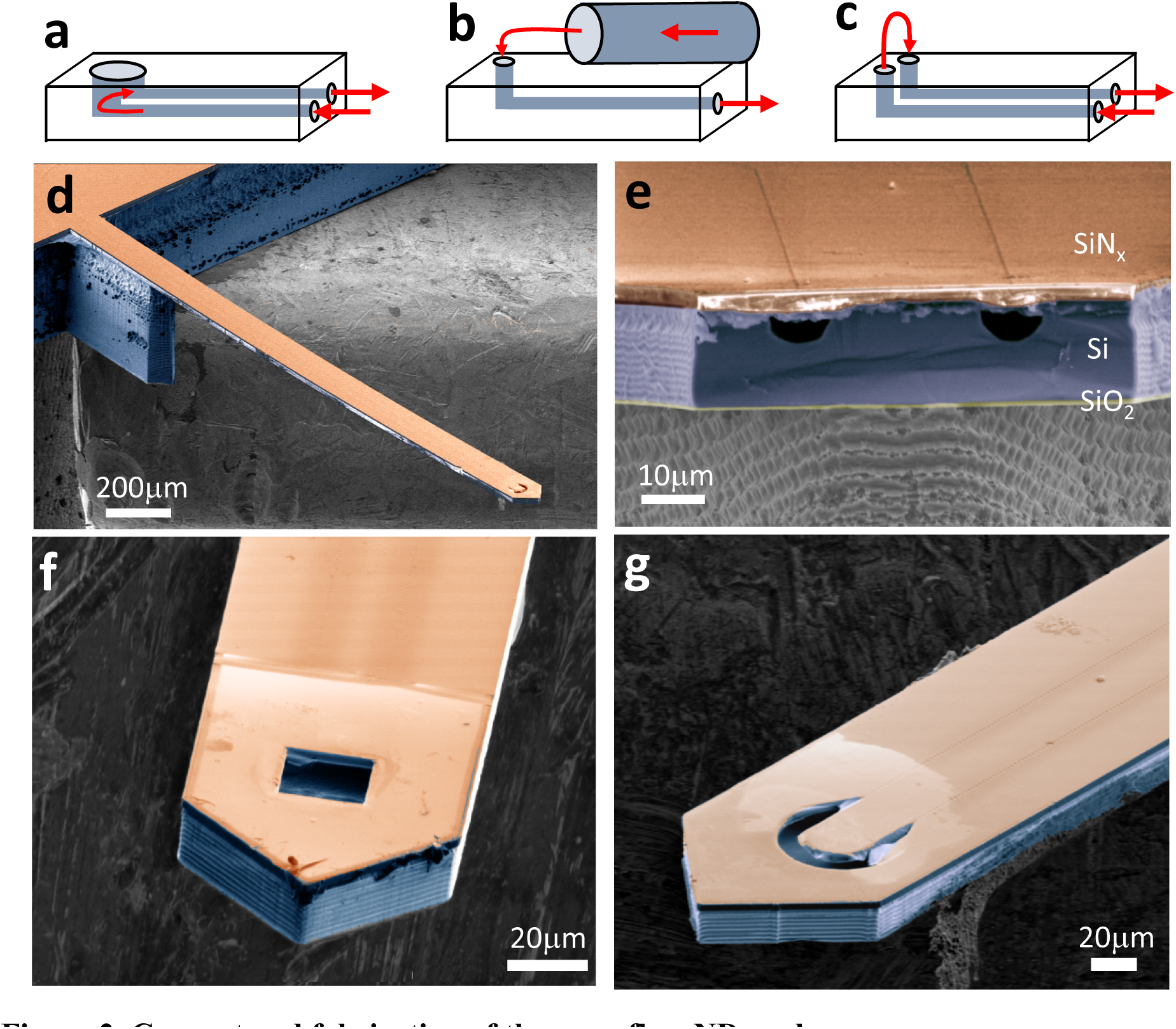
Concept and fabrication of the open flow ND probe. **A)** Schematics of in-plane open flow nanodialysis**. B)** Schematics of push-pull sampling with adjacent capillary of Ref. 10. **C)** Schematics of out-of-plane push-pull sampling of Ref.12. **D)** SEM image of the fabricated ND probe with 3mm long sampling needle. **E)** SEM cross-section of the probe needle showing 2 parallel microfluidic semicircular 5μm-radius channels sealed with a SiNx layer. **F)** SEM image of the tip of the completed open flow ND probe with the square sampling area design. **G)** SEM image of the tip of the completed open flow ND probe with the U-shape sampling area design.

The open-flow ND probe (Fig.2D) is fabricated following a silicon microfabrication process (Fig.S1A) that involves 5 layers of aligned photolithography, 2 thin film depositions and 10 etching steps. The probe sampling needle is designed to be several millimeters long to provide access to deep tissues. Fabrication on silicon-on-insulator (SOI) wafers with a 15μm device layer enables reduction of the probe needle cross-section to 17×70 μm^2^ (1260 μm^2^), which is the smallest demonstrated thus far (Fig.2E, Fig.S1B), and is approximately just a few cell bodies across thus minimizing tissue trauma and inflammation. This cross-section can accommodate up to 3 microfluidic semi-circular channels of 5μm radius etched into the Si device layer and sealed with a 3 μm thick SiNx layer (only 2 channels are used for the design in Fig.2E, Fig.S1B).

Two separate designs of the probe sampling area at the tip of the needle are explored: a 1540 μm^2^ square design (Fig.2F) and a more complex U-shape design with an area of 1720 μm^2^ (Fig.2G) with a gradual tapering of the channel width to maintain an in-plane laminar flow (see also Fig.S6).

### Realization of the in-plane pressure-balanced open-flow ND probe

To characterize the flows and recovery rates of the fabricated ND probe, the probe is released from the processed SOI wafer (Fig.S1CD,E) and packaged to provide fluidic interfaces with external microfluidic pumps (Fig.3A). Packaging involves attaching the chip to a 3D-printed holder, insertion of 3 microchannel stubs at the chip edge into glass capillaries, and sealing the junctions with UV-curable resin (Fig.S1F,G,H).

**Figure 3.**
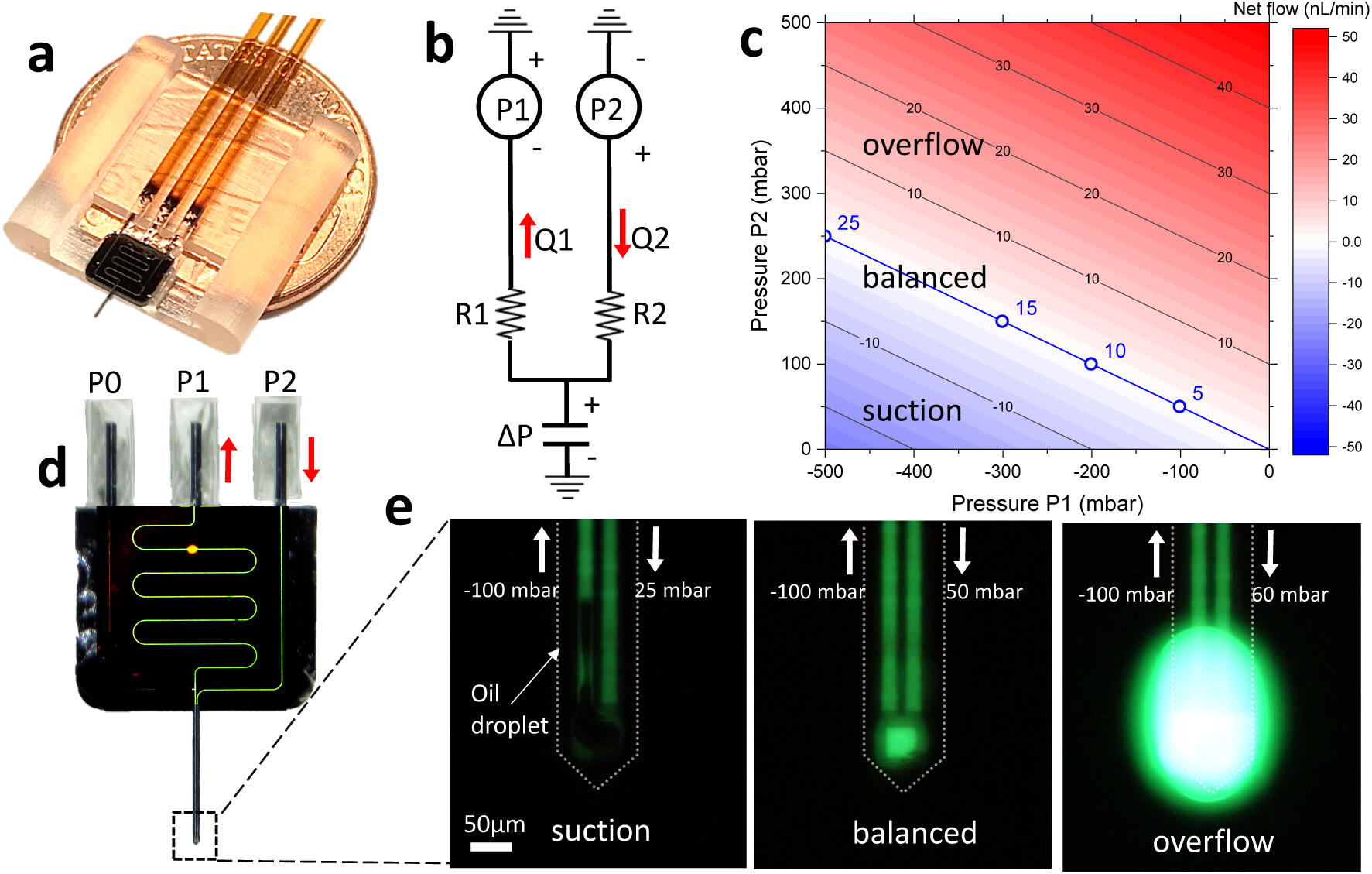
Realization of the in-plane pressure-balanced open-flow ND probe. **A)** Photograph of a packaged ND probe**. B)** Equivalent circuit diagram of pull (1) and push (2) arms of the microfluidic circuit. **C)** Calculated net flow rate as a function of a pressure balance between pull and push channels. Blue numbers correspond to flow rate at balanced pressure. **D)** Fluorescence microscope image of the packaged probe visualizing the inlet and outlet aqueous flows with dissolved 1mM fluorescein. **E)** Fluorescence microscope image of the sampling tip of the ND probe with the square-shape open sampling area under negative (left, oil suction), nearly zero (middle, balanced), and positive (right, dye overflow) differential pressures 𝛥 (SM Movie 1).

First, we assess the concept of the in-plane balanced open-flow sampling (Fig.3). The probe in this configuration can be analyzed as an equivalent circuit (Fig.3B) with hydrodynamic resistances R1 and R2, and pressures P1 and P2, corresponding to the pull and push flows, respectively. Since we are aiming to maintain the pressure at the open sampling site to be equal to the atmospheric pressure (or the intracranial pressure for measurements in the brain), we assume both circuit arms are grounded at the sampling site. The interface between the perfusate and the tissue can be elastic, hence we consider it to be a hydrodynamic capacitor that reflects the volume of the open sampling area changing with differential pressure 𝛥 = (𝛥_1_𝑅_2_ + 𝛥_2_𝑅_1_)⁄(𝑅_1_ + 𝑅_2_) applied to it. Hydrodynamic resistances R1 and R2 were estimated from the cross-section and lengths of the channels, and the measured flow velocity for a given pressure (Methods). These hydrodynamic resistances were used to predict the net flow rate at the open-flow sampling site (Fig.3C). Neglecting the capacitance, these flows are perfectly balanced (net flow is zero) when the differential pressure ΔP is zero corresponding to the blue diagonal in Fig.3C. Along this diagonal the in-plane flow rate can be tuned continuously from below 5nL/min on the lower right corner to above 25nL/min on the upper left corner.

To visualize and measure balanced flows, the packaged ND probes are attached to a glass slide with their sampling needle embedded into a 0.5mL reservoir (Fig.S2) filled with fluorinated oil (FC40, Sigma-Aldrich). Various pressures are applied to the chip’s push and pull channels (Fig.3D). Since the on-chip microfluidic channels are buried under an optically transparent SiNx channel cover, the standard fluorescence microscopy is used to visualize (Fig.3D) aqueous microfluidic flows (fluorescein in DI water) inside the buried channels. Figure 3E shows a series of frames from a movie (SM Movie 1) recorded during a sweep of 𝛥2 from 25mbar to 60mbar while keeping 𝛥1 constant at -100mbar (vertical cut in Fig.3C). When 𝛥 at the open-flow area is negative (Fig.3E, left) the oil is suctioned into the outlet channel forming a black elongated droplet, while a positive 𝛥 results in an overflow of dye solution (Fig.3E, right). A balanced flow is achieved when the push and pull flow rates are nearly equal, resulting in purely in-plane flow (Fig. 3E, middle) around the 𝛥 = 0 predicted from Fig.3C with margins ±30mbar (SM Movie 1). This margin is consistent with the expected Laplace pressure at the water/oil interface inside the open-flow area, which when overcome, results in suction (negative 𝛥) or overflow (positive 𝛥). .

### Characterization of the open-flow ND probe at balanced pressure

The established boundaries of the pressure balance correspond to a laminar flow of the perfusate in the open-flow sampling area with minimized out-of-plane bulk flow (Fig.4A). To measure the 𝑅 at room temperature (RT) the reservoir is filled with a solution of 1mM fluorescein in DI water and the the dye concentration in the dialysate channel is quantified by measuring the fluorescence intensity with a photomultiplier tube (PMT) attached to a fluorescent microscope (Fig.S2). For the square-shape open-flow area design the 𝑅 rapidly increases with decrease of the flow rate reaching (45.3% ± 6.3%) (mean ± standard deviation (SD), n=4) at 5nL/min (Fig.4B, black circles). The U-shape open area design exhibits a much higher 𝑅, reaching (83.3%±3.1%) (mean ± SD, n=4) at 5nL/min. The absolute flux of dye molecules across the open area interface is shown in Fig.4C. For the square-shape design (Fig.4C, black circles) it remains almost constant at (8819±1243) nmol/cm^2^/hour (mean ± SD, n=4) for all flow rates. However, for the U-shape design (Fig.4C, red circles) the flux exhibits a strong 4X increase from (14,204 ± 530) nmol/cm^2^/hour (mean ± SD, n=4) at 5nL/min to (61,030±3,525) nmol/cm^2^/hour (mean ± SD, n=4) at 38nL/min.

**Figure 4.**
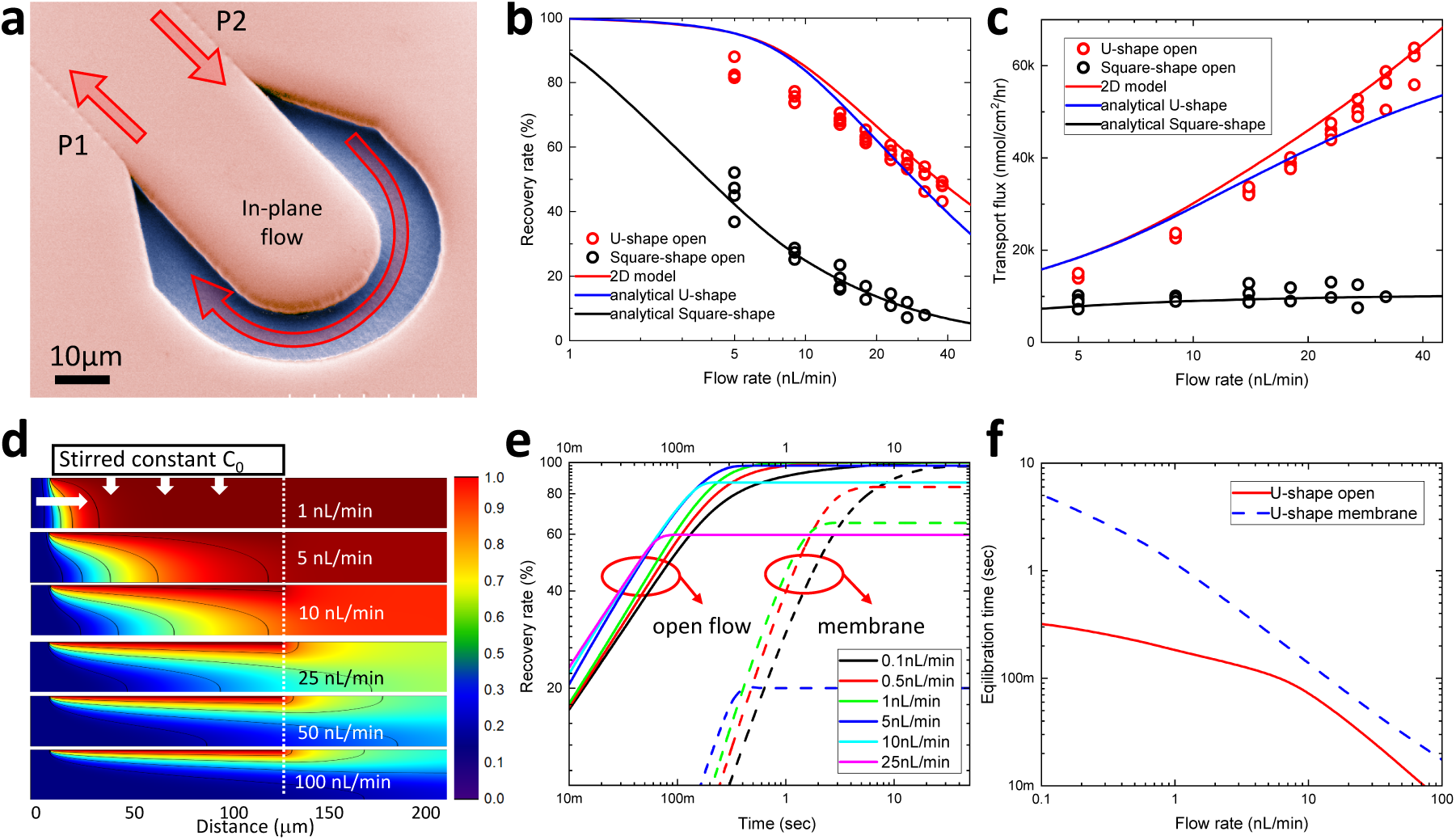
Characterization of the open-flow ND probe at balanced pressure. **A)** SEM image of the U-shape open sampling area**. B)** Recovery rates of the U-shape (red circles) and square-shape (black circles) open-flow ND probes measured at different pressure-balanced perfusate flows at RT. Red curve is the RR calculated by the 2D COMSOL model for 130μm length of the U-shape design. Blue and black curves correspond to fitting with Eq.1. **C)** Dye transport flux measured for the U-shape (red circles) and the square-shape (black circles) open-flow ND probes measured at different pressure-balanced perfusate flows. Red curve is the RR calculated by the 2D COMSOL model for 130μm length of the U-shape design. Blue and black curves correspond to fitting with Eq.1. **D)** Calculated 2D concentration profiles of in-plane dialysate flows. The colors represent concentration of fluorescein, with blue being 0mM and red being 1mM. **E)** Calculated transient recovery rates for different flow rates. Solid and dashed lines correspond to U-shape design with open area and integrated resistive membrane, respectively. **F)** Dependence of the equilibration time on flow rates for U-shape design with open area (solid red) and integrated resistive membrane (dashed blue).

To get further insight we performed numerical calculations using a 2D COMSOL model of a steady-state diffusive flux through an open sampling area with the channel depth of 10μm (Fig.S6) for different in-plane laminar flow rates (Fig.4D). Calculated concentration profiles can be understood by comparing the diffusion time 𝜏_𝐷_ it takes molecules to diffuse vertically to the bottom of the channel, to the residence time 𝜏_𝑅_ required for molecules to flow along the full length of the open sampling area of 130μm (Fig.S6). Since at low flow rates of a few nL/min, 𝜏_𝐷_ is shorter than or comparable to 𝜏_𝑅_, a nearly constant concentration profile is formed downstream across the whole channel depth (Fig.4D, top) with 𝑅 approaching 100% (Fig.4B, red). At high flow rates 𝜏_𝑅_ is much smaller than 𝜏_𝐷_, which leaves the bottom of the channel almost completely depleted of the analytes (Fig.4D, bottom) and therefore significantly diluting the dialysate, with a corresponding strong decrease of the 𝑅 (Fig.4B, red). The 2D numerical model produces results consistent with direct fitting using Eq.1 (Fig.4B, blue) assuming the transport resistance 𝑅_𝑑_ of 5.4 · 10^5^ sec/cm for an open area 𝑆 of 1760μm^2^. The 𝑅_𝑑_ is the largest at slow flows since the concentration gradient across the interface is the smallest, which results in diminishing flux also observed in the results of the U-shape design in Fig.4C. At higher flow rates the concentration gradient rapidly increases resulting in smaller 𝑅_𝑑_ and an increase of the flux. For the square design with similar area 𝑆 of 1540μm^2^, the 𝑅_𝑑_ is 6.5X larger reaching 3.5 · 10^6^ sec/cm (black curve in Fig.4B, C). Correspondingly, the flux is significantly reduced indicating a much shorter effective sampling length compared to U-shape design. Since the 𝑅 for the U-shape design is markedly better than that of the square-shape design, it is used exclusively for further experiments involving the ND probe with integrated nanoporous membrane.

Integration of a semipermeable membrane is expected to increase the transport resistance, which can potentially result in an increasingly long time for the concentration to reach the equilibrium steady-state gradients of Fig.4D thus degrading temporal resolution. To evaluate the equilibration time 𝜏_𝐸_ (time to reach 80% of the equilibrium steady state 𝑅) we used the transient 2D COMSOL model for different transport resistances (Fig.4E, solid lines). For an open area U-shape design (solid lines in Fig.4E) the calculated 𝜏_𝐸_ is below 1sec even at flow rate as small as 0.1nL/min. Corresponding dependence of 𝜏_𝐸_ versus flow rate is shown in Fig.4F, red. When the resistance is increased to 10^7^ sec/cm, the transient 𝑅 exhibits much slower dynamics (Fig.4E, dotted lines) with calculated 𝜏_𝐸_ reaching 1sec at 1nL/min (Fig.4F, blue).

### Characterization and integration of ultrathin Si nanoporous membrane

For our ND probe, we adopted a silicon nanoporous membrane^22^ (SiMPore, NY) (Fig.5A) that has been explored previously for hemodialysis^21^ and showed high permeability^25^ and molecular selectivity^26^. The membrane thickness is just (30 ± 2) nm (mean ± SD, n=5) with an average pore’s radius of (23.3 ± 6.6) nm (mean ± SD, n=208) with measured porosity of (15.5 ± 6.6) % (mean ± SD, n=10) (Fig.S4). The fluorescein transport flux of (2511 ± 502) nmol/cm^2^/hour (mean ± SD, n=3) through the Si membrane (Fig.5B) measured at 50°C using a side-by-side diffusion cell (Fig.S3) is 140X higher than measured for the reference AAO membrane (18 ± 3) nmol/cm^2^/hour (mean ± SD, n=3) with similar pore size and porosity, but with thickness of 50μm (InRedox, CO). While for the AAO membrane there is a significant selectivity between anionic fluorescein (332 Da) and methyl orange (327 Da) dyes, and cationic methylene blue (320 Da) dye, of similar molecular weights and hydrodynamic diameters (<0.5nm), the molecular flux through the ultrathin Si nanoporous membrane is independent of the molecules charge (Fig.5B, Fig.S4) since it is mostly defined by the pore discovery time rather than the transmembrane transport^26^.

**Figure 5.**
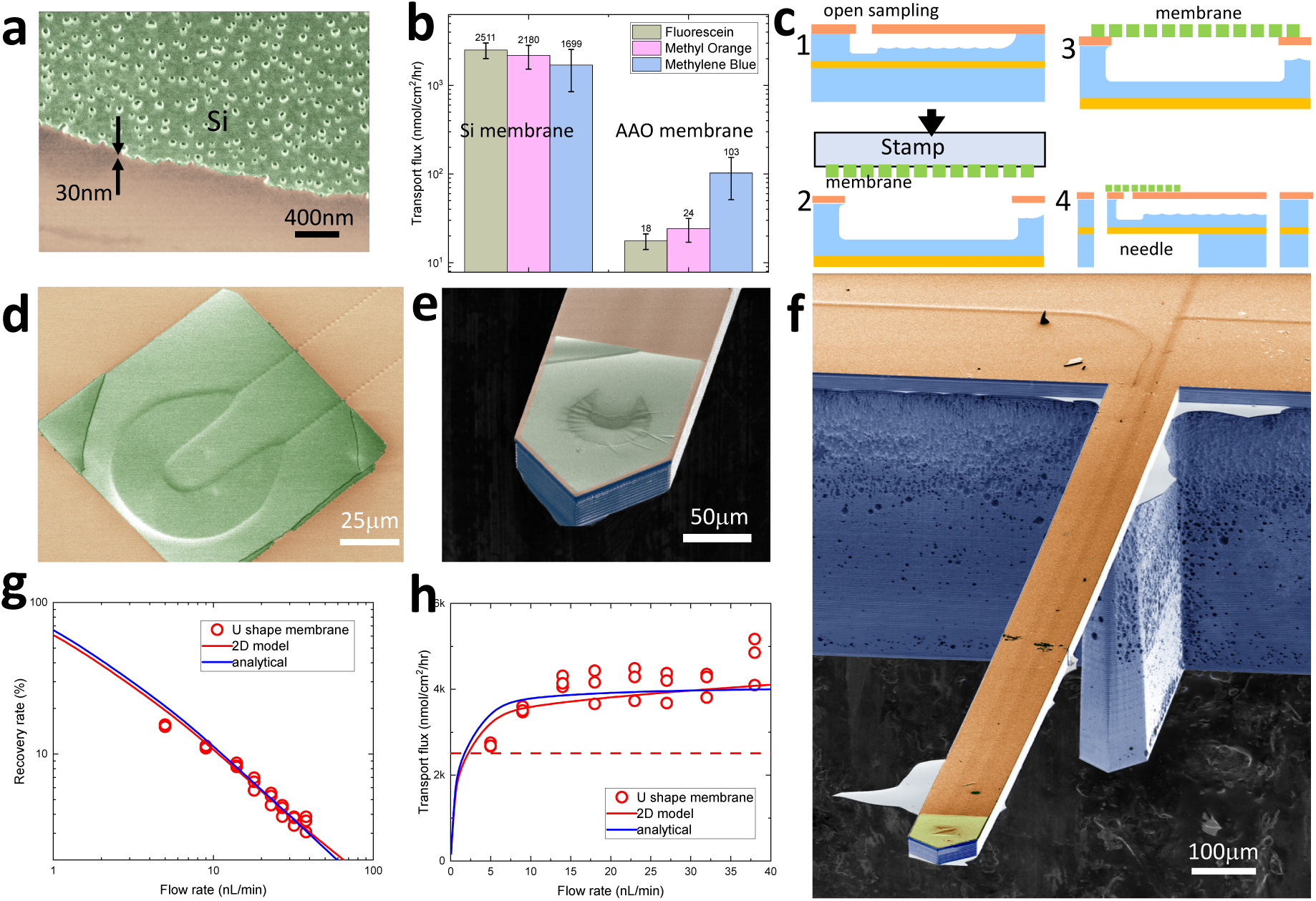
Characterization and integration of ultrathin Si nanoporous membrane. **A)** SEM image of the nanoporous ultrathin silicon membrane**. B)** Dye transport fluxes measured with nanoporous Si membrane and with AAO membrane for different dye molecules at 50°C. **C)** Fabrication steps to integrate the ultrathin membrane onto the open sampling area of the ND probe (step 1) using stamping technique (step 2, see also SM Movie 2), cleaning the wafer (step 3) and releasing the probe from the wafer with front and backside etching (step 4). **D)** SEM image of the membrane transferred on top of a U-turn sampling area. **E)** SEM image of the tip of the completed ND probe with a U-shape sampling area design. **F)** SEM image of the fully processed ND probe with integrated ultrathin Si nanoporous membrane. **G)** Relative recovery of a U-shape ND probe with a Si nanoporous membrane measured at RT. **H)** Dye transport flux of a U-shape ND probe with a Si nanoporous membrane at RT. Dashed horizontal line is the transport flux for a stand-alone silicon nanoporous membrane at 50°C.

To integrate the Si nanoporous membrane into the ND probe we developed a wafer-scale fabrication flow (Fig.5C, Fig.S5, SM Movie 2) adapting a stamping technique^27^ to detach the membrane from the host substrate using a polydimethylsiloxane (PDMS) substrate, align and attach it to the ND open sampling area (step 2 in Fig.5C), stamp and release the membrane (step 3 in Fig.5C), and clean the residue (Fig.5D). Once the membrane is in place, the wafer undergoes two deep reactive ion etching steps to undercut the sampling needle (step 4 in Fig.5C and Fig.5E). The released ND probe with a Si nanoporous membrane integrated onto the sampling needle with the U-shape design is shown in Fig.5F.

To measure the recovery rate (Fig.5G) and the dye transport flux (Fig.5H) of the ND probe, the sampling needle is immersed into a reservoir filled with 1mM fluorescein DI water solution at RT (Fig.S2A). While the flow rate dependencies (Fig.5G,H) are qualitatively similar to those observed with the open flow ND probe (Fig.4B,C), the 𝑅 at 5nL/min flow rate is decreased to (15.3 ± 0.3) % (mean ± SD, n=3) due to limited membrane porosity. The transport flux is (2610 ± 44) nmol/cm^2^/hour (mean ± SD, n=3) at slowest flow rate of 5nL/min that is close to that measured for the stand-alone membrane (dashed line in Fig.5H) with no flow. The transport flux increases to (4705 ± 550) nmol/cm^2^/hour (mean ± SD, n=3) at 38nL/min, which is among the highest reported for nanoporous membranes^22,23^. Both 2D numerical COMSOL model (red curves in Fig.5G,H) and analytical Eq.1 model (blue curves in Fig.5G,H) provide good fit to the flow rate dependencies in assumption of the total transport resistance 𝑅_𝑑_ + 𝑅_𝑚_ of 9 · 10^6^ sec/cm.

### On-chip segmentation of dialysate flow and on-chip droplet storage

Although the equilibration time for the ND probe with integrated Si membrane at 5nL/min flow rate is below 1 sec (Fig.4F), the overall temporal resolution for the ND probe can be significantly degraded by the lateral Taylor diffusion which occurs in the microfluidic channel that transports the collected dialysate for subsequent chemical analysis. Segmentation of the dialysate flow into a series of oil-isolated nL-volume droplets^28^ was shown to halt this diffusion and helped to reach temporal resolution in the range of just a few seconds^8,10–12,14^. The drastic reduction of channel cross-section down to 50μm^2^ enabled by our silicon platform and its operation at exceedingly slow nL/min flow rates allows for stable on-chip generation of droplets with volumes as small as just a few pL^24^. On-chip delivery of such droplets to online mass spectrometry (MS) analysis via an integrated nano-electrospray ionization (nano-ESI) nozzle^29^ or printing them one-by-one for matrix assisted laser desorption ionization (MALDI) MS^30^, we have previously demonstrated limits of detection (LOD) at the level of just a few attomoles^31^. However, such drastic reduction of flow rates and analyte volumes imposes a specific challenge of interfacing the segmented flows at the ND chip edge. Since the 27μm diameter of a 10pL droplet is much smaller than the inner diameter of a typical glass capillaries used to connect the chip to the external pumps, to store the droplets^32^, and to transport them to chemical analysis, droplets typically tend to intermix or coalesce^10^ thus significantly decreasing temporal resolution.

To address this problem, we designed an on-chip storage delay line (Fig.6) where collected droplets can be kept isolated from each other while preserving their order prior to hyphenation to MS analysis. To decrease the total hydrodynamic resistance 𝑅1 of the delay line (Fig.6A) its channel cross-section is increased to 357μm^2^ by increasing the channel width to 40μm. Oil flow controlled by the pressure 𝛥0 (Fig.6A) is transported through a channel 𝑅0 to the T-junction which operates in the squeezing regime (Fig.6B, SM Movie 3) to segment continuous aqueous flow into individual monodisperse (2.5% relative SD) aqueous plugs separated by oil droplets. The volume of the compartments can be continuously tuned (Fig.6C) from 40pL to 130pL by adjusting the oil-port pressure 𝛥0 and holding 𝛥1 constant (Fig.6D, SM Movie 4). The 26mm long delay line of Fig.6D can store anywhere from 20 (Fig.6D. left) to 60 (Fig.6D, right) aqueous samples corresponding to up to a minute of sampling time. The on-chip storage capacity is defined by the sampling frequency, droplet volume, channel length and hydrodynamic resistances, and therefore can be designed beforehand to meet experimental needs.

**Figure 6.**
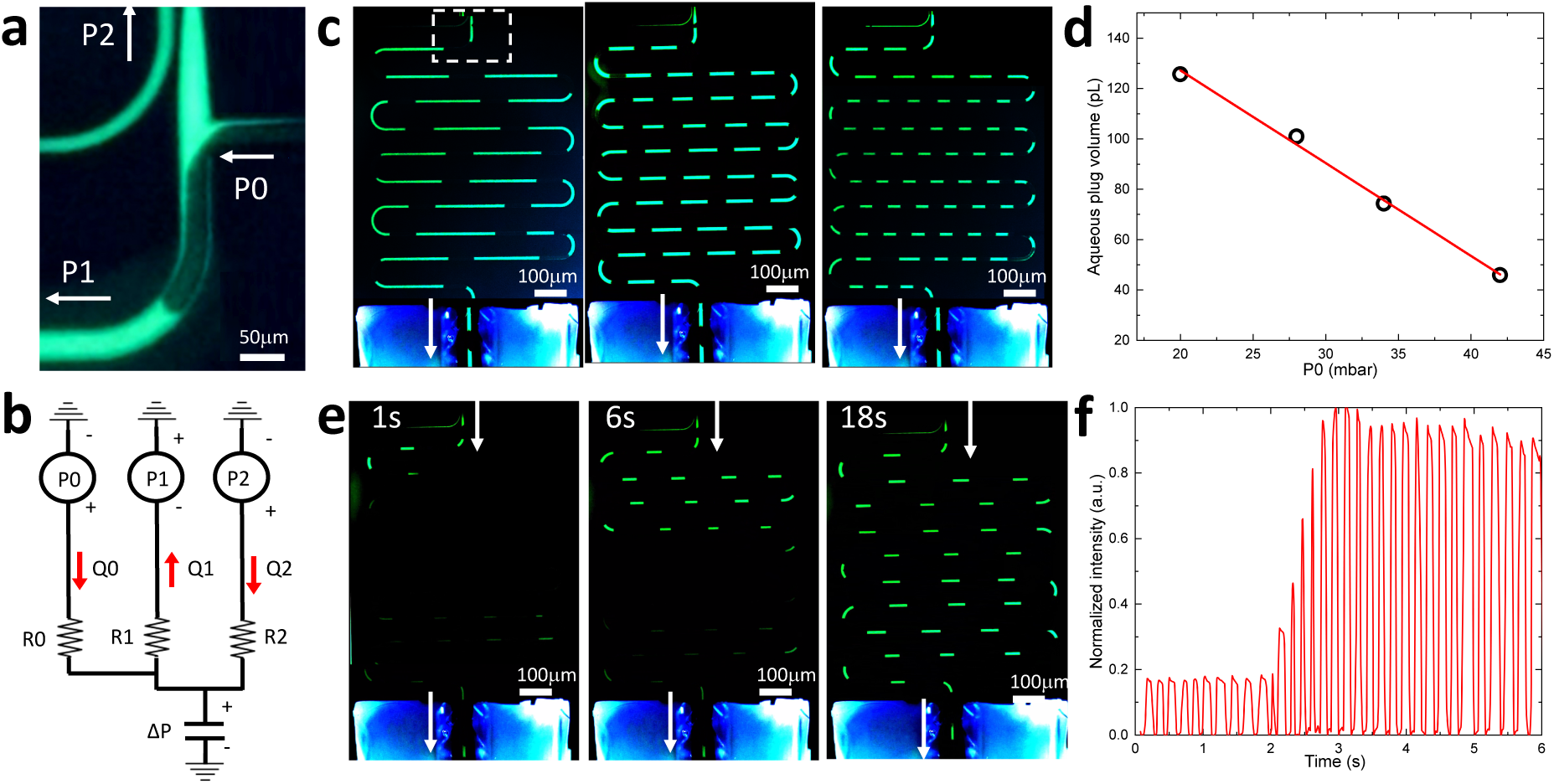
On-chip segmentation of dialysate flow and on-chip droplet storage. **A)** Fluorescence microscope image of the T-junction area (SM Movie 3). **B)** Equivalent circuit diagram with oil port P0 and a T-junction**. C)** Segmentation of a dialysate flow into compartments of different volume demonstrating a storage capacity of 20, 46, and 66 droplets, left to right. The dashed white rectangle in the left image corresponds to the location of a T-junction in A). **D)** Dependence of the aqueous plug volume on the P0 pressure while P1 is kept constant. **E)** Series of still frames from a video (SM Movie 5) taken while the concentration of fluorescein in the reservoir is rapidly changed. **F)** Fluorescence intensity measured with a PMT downstream during a rapid change in fluorescein concentration.

The temporal response provided by the flow segmentation is tested in Fig.6E by rapidly changing the fluorescein concentration in the reservoir while monitoring the fluorescence intensity in the downstream channels (Fig.6E, SM Movie 5). Only a few droplets are required to resolve fast changes in the concentration with a 20% to 80% threshold corresponding to just 450ms (Fig.6F).

## Discussion

Our results demonstrate that the designed silicon ND probe with integrated nanoporous membrane can provide high recovery rates, sub-second temporal resolution, and highly localized sampling, simultaneously, within a cross-section that is just a few cell bodies across. These unprecedented performance metrics are a direct result of a 100X down-scaling of the microfluidic channel cross-section, a corresponding drastic 100X reduction of flow rates to exceedingly slow few nL/min flows, and integration of a nanometer-thin nanoporous membrane with high transport flux into the probe sampling area. These engineered improvements were made possible by adopting a wafer-scale silicon microfabrication platform that provides precise control over channel dimensions, a high Young’s modulus to withstand pressure-controlled flows, and optical transparency for flow visualization using standard fluorescence microscopy. Further integration of on-chip droplet segmentation and a delay line for droplet storage enables sampling and manipulation of analytes at volumes of just a few pL. Integration with previously demonstrated hyphenation of segmented dialysate flows to sensitive MS methods using nano-ESI^29^ or MALDI-MS^30^ will enable quantification of small molecules at the level of just a few attomoles^31^. The proposed and demonstrated scaling strategy (Fig.1B,C,D) can potentially be extended even further in miniaturizing the sampling area, while keeping high recovery rate and temporal resolution. With mature silicon microfabrication technology, it is relatively straightforward to reduce the channel cross-section, and hence control the pressure-driven flow rates below 1nL/min (Fig.1D). Integration of ultra-thin membranes with high transport flux in the ND probe with the in-plane sampling geometry (Fig.4A) can enable reduction of a sampling areas down to just 100μm^2^ (Fig.1C). A corresponding reduction of dialysate volumes to be manipulated and analyzed at a femtoliter range, as well as the required increase of chemical sensitivity to zeptomoles levels, are both within reach, especially in view of recent developments in nanofluidics^33^ and in chemical quantification of single subcellular organelles^34^. Equilibration times for a membrane-integrated ND probe will inevitably become slower (Fig.4E) approaching several minutes, however, can be significantly improved by decreasing the membrane transport resistance (Fig.1C), or by adopting the concept of an open-flow ND (Fig.4A).

## Acknowledgements

Research reported in this publication was supported in part by the NINDS of the NIH grants UF1NS107677 and RF1NS126061. Work of CKB was supported in part by the NIH grant 1F31NS126018-01A1.

## METHODS

### ND probe fabrication of microfluidic channels

The ND probes are fabricated using a silicon-on-insulator (SOI, UltraSil) wafer with a 425 µm handle thickness, a 1µm buried oxide (BOX) layer, and a 15µm device layer. First, a 300nm of SiNx is deposited using plasma enhanced chemical vapor deposition (PECVD) (PlasmaPro 100, Oxford Instruments) to serve as a hardmask for subsequent etching of the Si device layer (Fig. S1A.1). The SiNx hardmask is then patterned with rows of perforations using a direct laser lithography system (MLA 150 Heidelberg) followed by reactive ion etching (RIE) using an inductively coupled plasma (ICP) ICP-RIE etcher (PlasmaPro System100, Oxford Instruments), leaving perforations of 1µm diameter exposing the Si device layer below. Next, the exposed Si is isotropically etched using XeF2 gas (Xactix, SPTS). During etching the neighboring undercuts from each perforation are merged, forming a continuous microfluidic channel buried underneath the SiNx hardmask (Fig.S1A.2). The size of the perforations, distance between them, and the etching time control the channel cross-section that can be tuned from a semicircular 40 µm^2^ profile with 5µm radius (single line of perforations, Fig.S6E) up to 40×10µm shape with 360 µm^2^ cross-section (4 rows of perforations, Fig.S7). The perforations in the SiNx hardmask layer are then sealed by deposition of 3µm low stress SiNx via PECVD (Fig. S1A.3, Fig.S1B).

### ND probe open sampling area fabrication

The open sampling sites with square-shape or u-shape designs (Fig.2F,G) are defined in the same hardmask layer as the perforations. Therefore, the isotropic Si etching results in a deeper and wider profile than in the buried microfluidic channels (Fig.S6) that gradually changes from a 40 µm^2^ (Fig.S6E) to 260 µm^2^ (Fig.S6D) cross-section, effective slowing down the perfusate flow velocity to achieve higher recovery rates.

### ND probe perimeter definition and release

Lastly, to define the probe perimeter and to release the probe from the wafer, a series of 200 µm wide trenches are patterned on the device side using direct write lithography (Fig. S1A4). The exposed SiNx is etched using ICP-RIE, and the 15µm thick Si device layer is etched down to the BOX layer using deep reactive ion etching (DRIE) (LpX Pegasus, STS). The BOX layer is etched further down to the handle Si wafer using ICP-RIE. The backside-aligned lithography (EVG620, EVG) is repeated on the handle wafer side followed by a DRIE step to etch all the way through the 425µm thick handle wafer (Fig.S1C). The resulting probe is attached to the wafer via a set of narrow bridges that can be easily broken to release the probe with a sampling needle attached to the probe base (Fig.S1D) containing a buried network of microfluidic channels connecting sampling, droplet segmentation, and storage delay line components.

### ND probe packaging

The fabricated ND chip is released from the wafer and packaged to provide fluidic interfaces (Fig.S1E-H). To interface with external microfluidic pumps, the 170µm-wide by 425µm-thick silicon stubs on the chip base (Fig.S1F) are shaped to fit into 15cm-long 700 μm ID silica capillaries (TSP700850, Polymicro). The polyimide coating of the glass capillaries is burned out to allow transmission of UV light in subsequent steps. Precise alignment of all 3 capillaries simultaneously is achieved under a microscope with the aid of 3D printed holders designed for handling the probe and capliaries (Fig.S1G) and a pair of x-y-z micromanipulators. The connected stubs are sealed with UV curable resin (NOA 68T, Norland) to enable continous operation under high differential pressures (Fig.S1H).

### Integration of ultrathin nanoporous membrane

Integration of the nanoporous Si membrane into the process flow is introduced after the formation of the microfluidic channels and open sampling sites (Fig.5C) between steps 3 and 4 from Fig.S1A using a wafer-scale fabrication flow (Fig.5C, Fig.S5, SM Movie 2). The 30nm-thin silicon nanoporous membrane (US100-P30Q33, SimPore) with 500×500 µm^2^ surface area is detached from the Si host substrate using a polydimethylsiloxane (PDMS, Sylgard 184, DOW Chemical) stamp covered with a 2 µm-thin polymethyl methacrylate (PMMA, MicroChem) layer (Fig.S5A, step 1 SM Movie 2). The wafer is cleaned, treated with oxygen plasma, and pre-heated above the glass transition temperature of the PMMA to ease the attachment and release of the stamp from the wafer surface (Fig.S5B, step 2 SM Movie 2). The stamp with the attached membrane is aligned to several ND open sampling sites simultaneously (Fig.S5B) under a microscope and attached to the wafer (step 2 SM Movie 2). Then the PDMS stamp is lifted off and the wafer with attached membrane is cooled down to room temperature. PMMA residue is removed in acetone for 5 hours at 50 ℃. Successful attachment and detachment of the PMMA layer can be seen by the moving meniscus at the PMMA-substrate interface (red arrows in Fig. S5A,B and SM Movie 2). Even though the membrane is just 30nm thick and is hanging over a large sampling area, it is mechanically robust enough to withstand deposition of thick photoresist layers, double-sided lithography, long DRIE steps, photoresist stripping and cleaning steps (Fig.5E,F, Fig.S6).

### Molecular transport rate measurements in nanoporous membranes

To characterize the pores of the silicon nanoporous membrane and a reference anodic aluminum oxide (AAO) membrane (AAO-010-020-050, InRedox), SEM images were taken (Hitachi S-4800) at high magnification (Fig. S4A) in several areas of the membranes. The raw images were thresholded to isolate pore perimeters from the background, and pore radii, density, and porosity were measured using ImageJ (Fig. S4B). Prior to transport measurements, all membranes were exposed to oxygen plasma for 15 min.

To measure the diffusion transport rate, the membranes were placed in the middle of a side-by-side diffusion cell (5G-00-00-15-05, PermeGear) (Fig.S3A) with identical 5mL feed and permeate volumes to avoid hydrostatic built-up. To measure the transport of molecules with similar weight but different polarity through the membranes, 1mM solutions of fluorescein (Sigma-Aldrich), methyl orange (ACS reagent, dye content 85%, Sigma-Aldrich), and methylene blue (Sigma-Aldrich) in DI water (18MΩ, Millipore) were used to fill the feed cell. To ensure uniform concentration distributions in the feed and permeate cells, both cells were continuously stirred using magnetic stir bars. The diffusion cell was held at a constant 50°C temperature monitored with a thermo-pair. Optical transmission through a receiving permeate cell was analyzed (Fig.S3B) using a blue light from a fiber-coupled LED (490nm, M490F4, Thorlabs) collimated with a pair of high NA fiber optic collimators (F950FC-A, Thorlabs) and recorded using a fiber-coupled UV-VIS spectrometer (Flame Miniature Spectrometer, Ocean Optics). For uniform concentration fields the molar concentration *c* of the analyte in the permeate cell was calculated from the Beer–Lambert law:

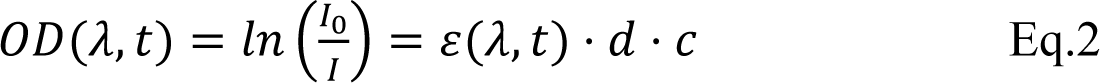

where *OD* is optical density, *ε* is the molar absorption coefficient, and *d* is the thickness of the cell. The transmission through the permeate cell was calibrated by measuring the transmission (Fig.S3C), calculating the optical density OD (Fig.S3D), and applying a linear fit of OD vs concentration (Fig.3E) for fluorescein (Fluo), methyl orange (MO), and methylene blue (MB) at wavelengths of 498nm, 503nm, and 473nm, respectively. Eq.1 produced a good fit of the Beer-Lambert law for ODs of at least up to 5 with R^2^ > 0.992 and Pearson rp > 0.995 for all analytes (Fig.S3E). Above the OD of 8 the dependence became nonlinear (Fig.S3D) and was not used fitting.

The temporal change of the OD also showed good linearity (R^2^ > 0.995, rp > 0.998) for all analytes (Fig.S4C) for long times, as the experiments lasted as long as several days. Similar dependencies were observed for the AAO membrane (Fig.4D) with high linearity for Fluo (R^2^=0.998, rp=0.999) and MB (R^2^=0.986, rp=0.993) with somewhat lower values for MO (R^2^=0.924, rp=0.961). The slopes of the linear fits of OD vs time (Fig.S4C) were normalized on corresponding calibration curves of OD vs concentration (Fig.S3E) to calculate temporal changes in concentration, transport resistances, and transport fluxes reported in Fig.5B.

### Visualization and characterization of microfluidic flows with fluorescence microscopy

To visualize microfluidic flows, the fully packaged ND probe (Fig.3A) was connected to external pressure-controlled pumps (Flow-EZ Module, LU-FEZ-7000 and LU-FEZ-N800, Fluigent) for push (P2 in Fig.6B) and pull (P1 in Fig.6B) aqueous flows, and a push oil flow (P0 in Fig.6B). Since the top SiNx layer of the probe is transparent for visible light, standard fluorescence microscopy was used. Imaging and characterization of flows and droplets was done with an inverted fluorescence microscope (IX73, Olympus) equipped with a photomultiplier tube (PMT 2101, Thorlabs), video cameras (EOS Rebel T7i, Canon, and Hero8, GoPro) and an LED lamp at 400nm wavelength (pE-300 White, CoolLED) for fluorescence excitation. To visualize the flows (e.g. Fig.3D,E), a fluorescein solution in DI water was used at various concentrations up to 1mM as specified for each experiment (e.g. Fig.3D).

### Calibration and measurements of diffusion fluxes in ND probes

To visualize the suction or oveflow of aqueous flows (Fig.3E), the fully packaged ND probe was attached to a glass slide with its sampling needle embedded into a 0.5mL reservoir (Fig.S2A) filled with fluorinated oil (FC40, Sigma-Aldrich), and various pressures were applied to the chip’s push (P2) and pull (P1) channels (Fig.3E, SM Movie 1).

To measure fluorescein concentration inside the microfluidic channels, the image of a microfluidic channel formed with a 40X objective of the inverted microscope with a field of view (FOV) of 50μm diameter (inset in Fig.S2B) was projected onto a PMT entrance slit, and the voltages were sampled and recorded with a digital multimeter (DMM6500, Keithley). Gain settings for the PMT and power settings for the excitation source were chosen to ensure the highest dynamic range. Example measurements of PMT voltages for 4 consecutive measurements of diffusive transport through an ND probe with integrated Si nanoporous membrane is shown in Fig.S2B. The flow rate was adjusted by changing the P1 and P2 pressures while keeping the pressure at the probe sampling area balanced (Fig.3C).

To convert the recorded PMT voltages (Fig.S2B) to concentrations, recovery rates (Fig.5G), and fluxes (Fig.5H), the calibration of PMT voltage was performed (Fig.S2C) by flowing solutions of known Fluo concentration at a constant 20nL/min flow rate resulting with a reasonable linear fit (R^2^=0.965, rp=0.975). To ensure that the flow rate itself did not affect the calibration, a set of measurements were performed for a constant concentration (1mM and 10μM) with flow rate varied (Fig.S2D).

### Flow segmentation and measurements of hydrodynamic resistance

To segment aqueous flow into a series of monodisperse plugs, oil flow controlled by the pressure P0 (Fig.6A) is transported through a channel R0 to the T-junction which operates in the squeezing regime (Fig.6B, SM Movie 3). The flow rate ratio between the aqueous and oil flows is controlled by precise balance of the P0 and P1 pressures (Fig.6D) that defines the volume and frequency of the droplets. As oil droplets are passing through a microscope FOV, the PMT voltage is strongly modulated enabling detection of the flow velocity. For segmented flows with relatively small oil droplets, the aqueous hydrodynamic resistance can be estimated by linear fitting of the measured flow velocity vs ΔP differential pressure using the known cross-section and length of the microfluidic channel. Normalized hydrodynamic resistances for aqueous (R1 and R2 in Fig.3B and Fig.6B) flows are measured to be 60 mbar/pL/sec/m and 574 mbar/pL/sec/m for FC40 oil (R0 in Fig.6B). These numbers are in close agreement with predictions using the analytical equation for a rectangular channel cross-section:

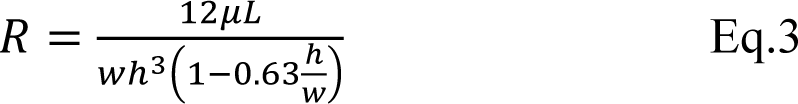

where 𝑤 and ℎ are the width and height of the channel, 𝐿𝐿 is the channel length, and 𝜇𝜇 is the dynamic viscosity taken as 8.9·10^-4^ Pa·sec for water and 4.1·10^-3^ Pa·sec for FC40 at RT.

### Measurements of temporal response

The temporal response of the ND probe with oil-segmented flow (Fig.6E) was tested by equilibrating the ND probe in a reservoir and then rapidly altering the fluorescein concentration surrounding the probe by using a pipette filled with 1mM Fluo solution while monitoring fluorescence of aqueous plugs downstream (Fig.6F). Droplet volumes were estimated by measuring droplet lengths and multiplying them by the cross-sectional area of the channel.

### Numerical modeling of steady-state diffusion fluxes for open flow ND probe

Numerical calculations using a 2D model of diffusion flux through an open sampling area for different in-plane laminar flow rates (Fig.4D) were performed using COMSOL Multiphysics 6.1. The microfluidic channel within the sampling area was modeled as a rectangular cross-section with a 11µm height and 23µm width to mimic the cross-section observed in experiment (Fig.S6). The U-shape opening was modeled as a straight channel with a length of 130µm, close to the arc length of the U-shape opening (Fig.S6). Small sections at the inlet and outlet of the sampling area with 10µm and 100µm length, respectively, were added to the model to simulate regions within the push and pull channels, respectively. The fluid flow within the microfluidic channel was modeled with the Navier-Stokes equations by using the “Laminar Flow” module. Convective and diffusive transport of analytes was modeled with the “Transport of Diluted Species” module. The left-side and right-side boundaries were set as an inlet and outlet respectively, with a fully-developed flow at equal constant flow rates (1, 5, 10, 25, 50, and 100nL/min) and entrance thicknesses equal to the width of the sampling area. The top and bottom boundaries of the microfluidic channel, including the boundary between the sampling area and the external volume, were specified with a no-slip boundary condition that assumes no convection into/out of the sampling area into the outside volume, and that the flow velocity at the boundary is equal to zero. To model the diffusion of analyte into the probe, the top sampling area boundary with the external volume was specified with a constant concentration boundary condition of 1mM fluorescein, with fluorescein having a diffusion coefficient of 4.9×10^-6^ cm^2^/s in water at RT and close to neutral pH. An integrated total flux [mol/s] of the analyte was taken at the outlet boundary and used to calculate recovered concentration. For modeling the steady state equilibrium fluxes (Fig. 4B, C, D), a “Stationary Study” was used.

### Numerical modeling of steady-state diffusion fluxes for ND probe with nanoporous membrane

To model the diffusion flux through an integrated nanoporous membrane (Fig. 4C), the top sampling area boundary was replaced with a “Thin Diffusion Barrier” with a no-slip boundary condition. The membrane thickness is set to 30nm and membrane diffusion coefficient to 2×10^-9^ cm^2^/s, a value derived from the experimentally measured transport flux in Fig.5B. Since the constant concentration boundary condition no longer applied to the top sampling area boundary, an additional fluidic cell with length of 130µm and height of 1µm was added above the sampling area to simulate a well-stirred solution at a constant 1mM concentration of fluorescein at 100µL/min flow rate. All other boundaries of this additional cell were set to a constant concentration of 1mM fluorescein.

### Numerical modeling of transient diffusion fluxes

To model the transient diffusion process (Fig.4E,F) prior to settling into a steady-state equilibrium, a “Time-Dependent Study” was utilized. The initial concentration was set to 0mM for both the inlet microfluidic channel and within the sampling area. The concentration in the additional fluidic cell above the sampling area is changed stepwise to 1mM and the temporal evolution of the concentration profiles is calculated for the following 60 sec. To resolve short-time and long-time transients of concentration gradients, gradated time bins were used starting from 0.1ms for the 0 to 1ms interval, 0.5ms bin for the 1ms to 332ms interval, and a 50ms bin for the rest.

## Supplementary movies

***SI Movie 1. Balanced in-plane flow.***

Fluorescence microscope video of the sampling tip of the ND probe with the square-shape open sampling area. Video is recorded while the pull pressure P1 is kept constant at -500 mbar and push pressure P2 is adjusted between 0 mbar and 360 mbar. When the differential pressure 𝛥 at the tip end defined by the hydrodynamic circuit of Fig. 3B is negative the oil suction is observed. At positive differential pressures dye overflow is observed. Balanced flow corresponds to nearly zero differential pressures with P1 between 200 mbar and 320 mbar in agreement with Fig.3C. Still frames from a similar movie are used in Fig.3E.

***SI Movie 2. Integration of Si nanoporous membrane into ND probe.***

The 30nm-thin silicon nanoporous membrane with 500×500 µm2 surface area is detached from the Si host substrate using a PDMS stamp covered with a 2 µm-thin PMMA layer. The wafer is cleaned, treated with oxygen plasma, and pre-heated above the glass transition temperature of the PMMA. The stamp with the attached membrane is aligned to several ND open sampling sites simultaneously under a microscope and attached to the wafer. Then the PDMS stamp is lifted off and the wafer with attached membrane is cooled down to room temperature. Successful attachment and detachment of the PMMA layer can be seen by the moving meniscus at the PMMA-substrate interface. Still frames from a similar movie are used in Fig.S5.

***SI Movie 3. On-chip dialysate segmentation into pL-volume droplets.***

Fluorescence microscope movie of the T-junction area showing segmentation of the dialysate flow into individual isolated compartments separated by FC40 oil droplets. Segmentation operates in the squeezing regime producing highly monodisperse droplets. Still frame from this movie is used in Fig.6A.

***SI Movie 4. On-chip storage of pL-volume droplets.***

Fluorescence microscope movie of the delay line area of the chip showing segmentation of the dialysate flow into monodisperse plugs with volume and frequency defined by the differential pressure 𝛥 between P1 and P2, as well as by P0 pressure which defines the flow rate ratio between aqueous and oil flows. Still frames used in Fig.6E are taken from similar movies recorded at different pressures.

***SI Movie 5. Temporal response.***

Fluorescence microscope movie of the delay line area of the chip recorded while the concentration of fluorescein in the reservoir rapidly increased. Still frames from this movie are used in Fig.6E.

**Fig S1.**
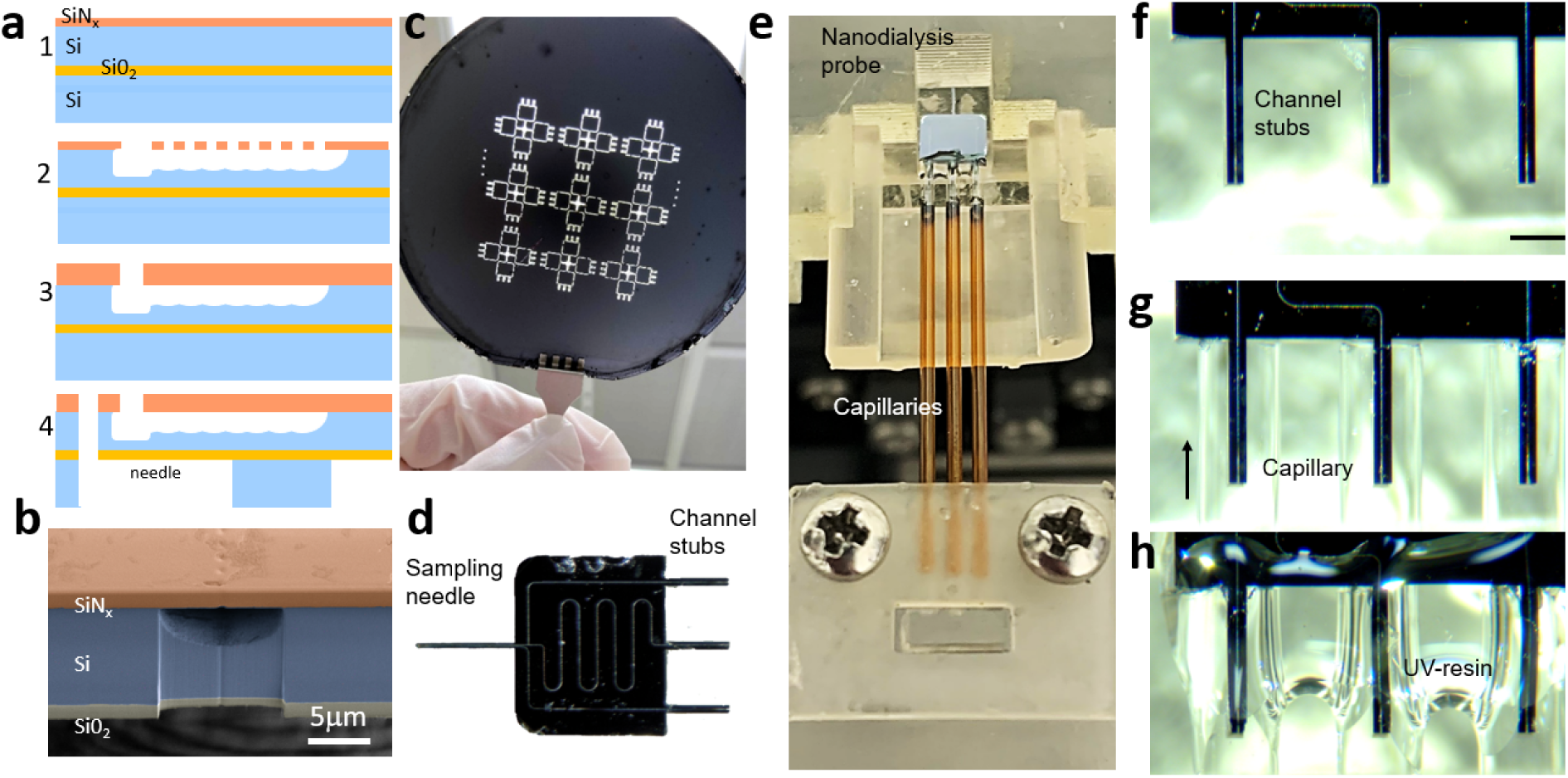
Fabrication and packaging of ND probe. **A)** Cross-section schematic of fabrication steps including (1) SiNx hard mask deposition on SOI wafer. (2) Channel formation by isotropic XeF2 etching of silicon through perforations in SiNx hardmask layer. (3) channel sealing with SiNx deposition, (4) ND probe perimeter and needle undercut defined by a double-side DRIE. **B)** SEM cross-section of the ND probe needle showing buried microlfuidic channel. **C)** Photograph of processed wafer with etched through perimeter trenches. **D)** Photograph of the released ND probe with a sampling needle, a probe base with 3 channels, T-junction, delay line, and packaging stubs. **E)** Packaging of the ND probe with 3 glass capillaries. **F)** Microphotograph of 3 stubs at the edge of the ND probe base. **G)** Microphotograph of 3 stubs aligned and inserted into cappolaries. **H)** Microphotograph of fully packaged ND probe with UV-resin sealed junctions.

**Fig S2.**
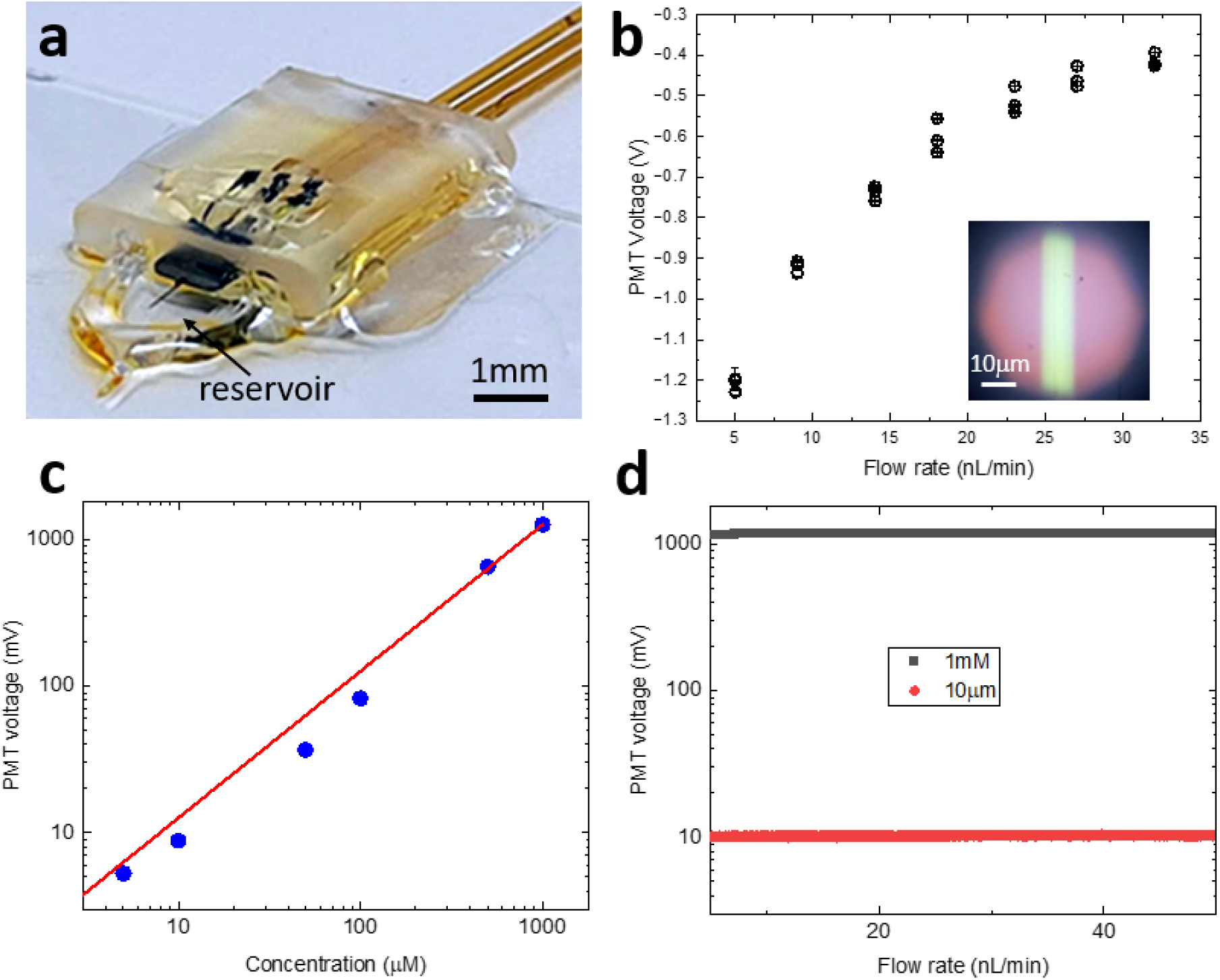
Calibration and measurements of concentration dependence. **A)** Photograph of a fully packaged ND probe attached to the glass slide with the sampling needle immersed into the reservoir. **B)** Inset: microphotograph of a channel filled with fluorescein solution that is used to measure flurescence intensity with PMT. Main: PMT voltages measured with 4 consecutive sweeps of the pressure-balanced flow rate while sampling from the resevoir containing 1mM of fluorescein using ND probe with integrated nanoprous membrane. **C)** Calibration of PMT voltage vs concentration measured at constant flow rate. **D)** Invariance of flurescence intensity vs flow rate for constant concentration.

**Fig S3.**
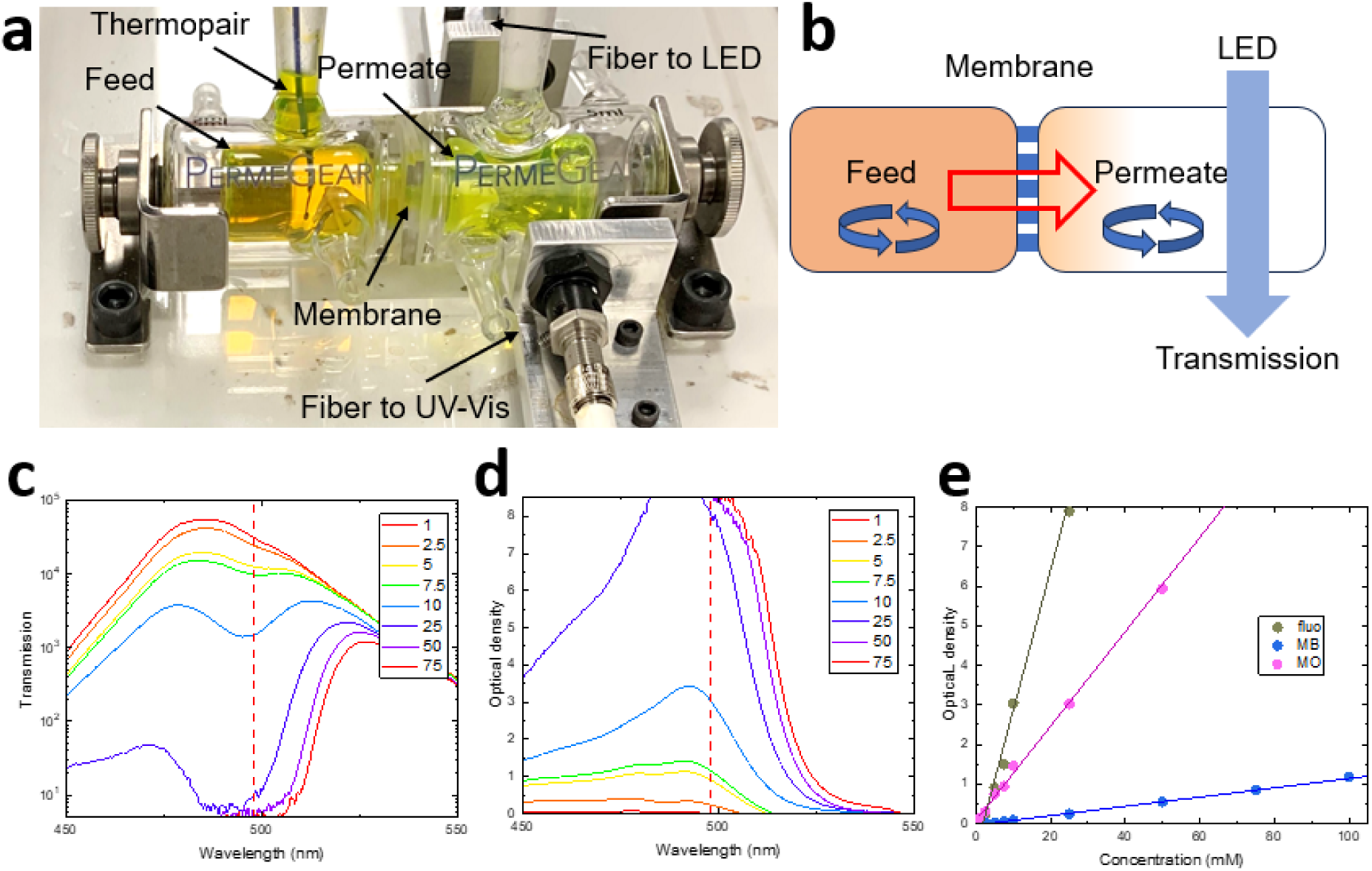
Calibration and measurements of membrane diffusive transport. **A)** Photograph of a side-by-side diffusion cell with fiber-coupled transmission measurement apparatus. **B)** Schematics of the setup in A) showing membrane separating the feed and permeate cells, magnetic stirrers, and fiber-coupled transmission apparatus. **C)** Transmission spectra measured through permeate cell for different concentrations of fluorescein in DI water. **D)** Optical density spectra calcuated from C). **E)** Calibration curves obtained from D) for fluorescein (black), methylene blue (blue), and methyl orange (magenta).

**Fig S4.**
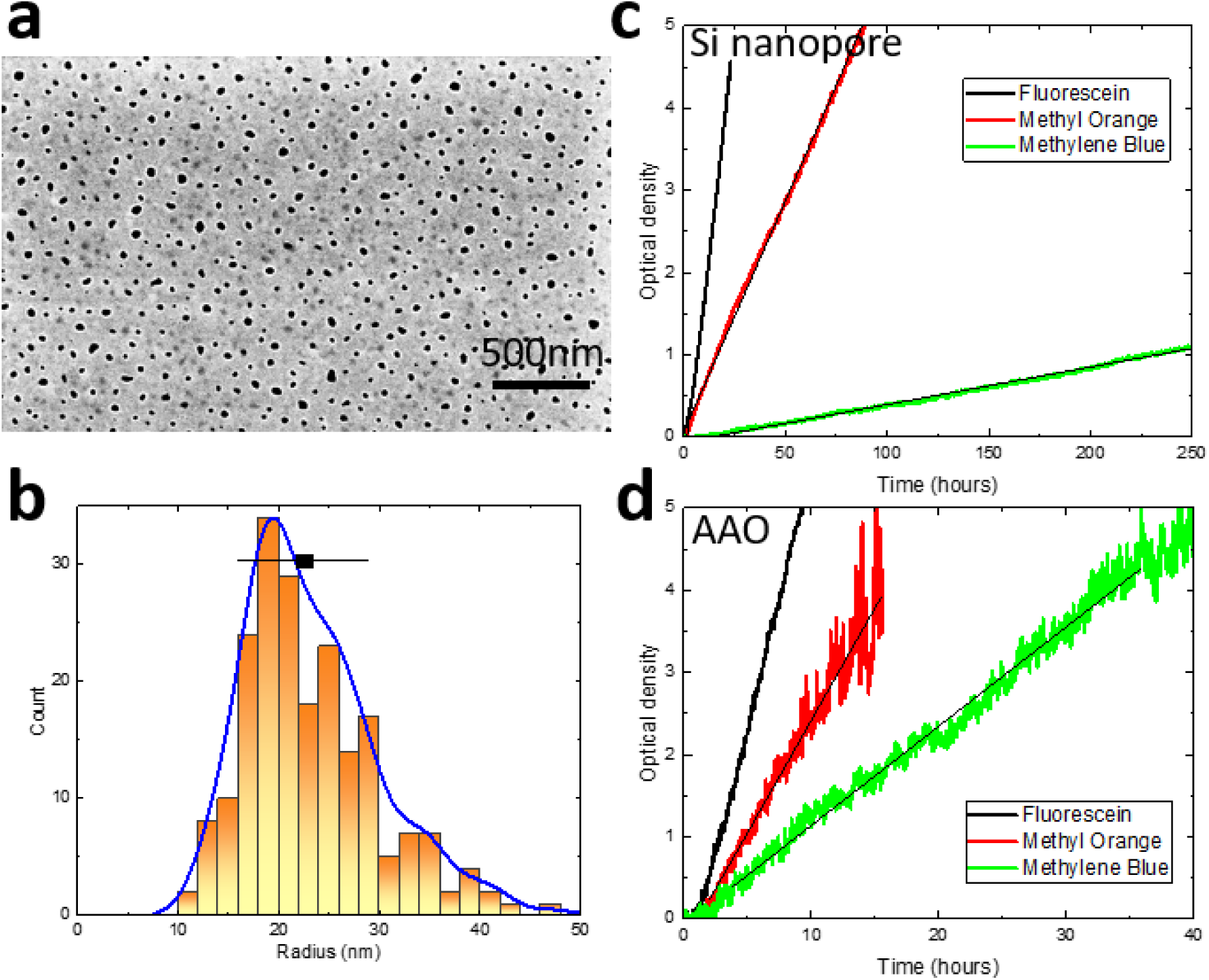
Porosity of Si nanopore membrane and diffusion measurements. **A)** SEM photograph of the ultrathin nanoporous membrane. **B)** Histogram of the pore radii measured from a set of SEM photographs as A). Box and whiskers correspond to median and standard deviation. **C)** Measurements of diffusive transport through an ultrathin Si nanoporous membrane. Temporal dependence of the optical density in the permeate cell while the feed cell is loaded with 1mM solution of fluorescein (black), methyl orange (red), or methylene blue (green). **D)** Same as C) for AAO membrane with 50μm thickness.

**Fig S5.**
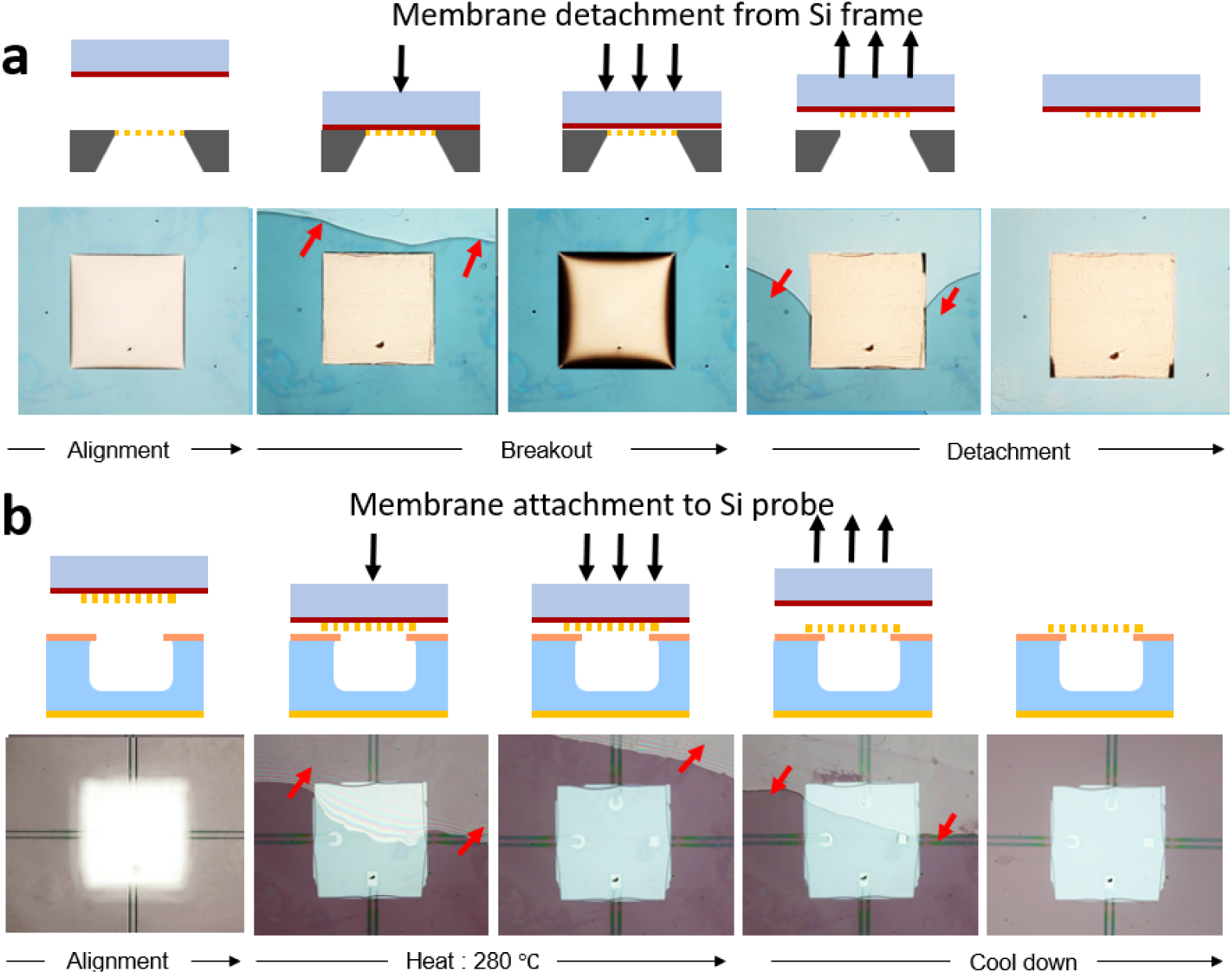
Integration of ultra-thin nanoporous Si membrane onto ND probe. **A)** Consecutive steps of membrane attachment to PDMS/PMMA stamp, breakout off the host Si chip, and detachment of the stamp. Red arrows show meniscus sweeping during membrane attachment and detachment. **B)** Consecutive steps of stamp alignment to the target wafer with ND probes, its attachment to the wafer surface and subsequent release. Red arrows show meniscus sweeping during membrane attachment and detachment.

**Fig S6.**
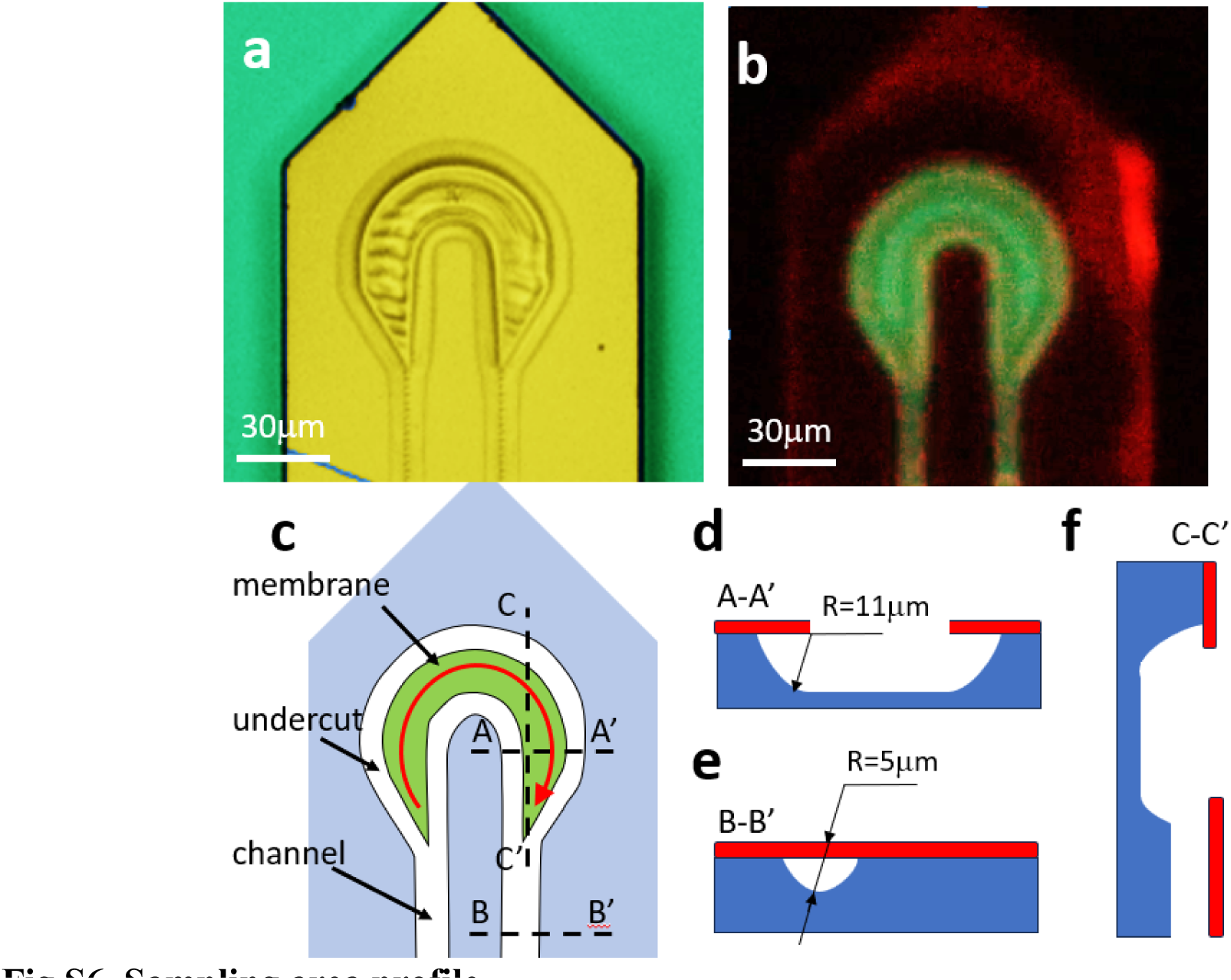
Sampling area profile. **A)** Optical microphotograph of the sampling area with integrated nanoporous membrane. **B)** Fluorescence microphotograph of the sampling area with integrated nanoporous membrane. Green color is fluorescence of fluorescein flowing through channels. **C)** Schematics of the sampling area layout showing channels, membrane area, and sampling area undercut. **D)** Schematic of the layout and cross-sections in the sampling area. **E)** Schematic cross-section in the channel area **F)** Schematic cross-section along the tapered connection between the two areas.

**Fig S7.**
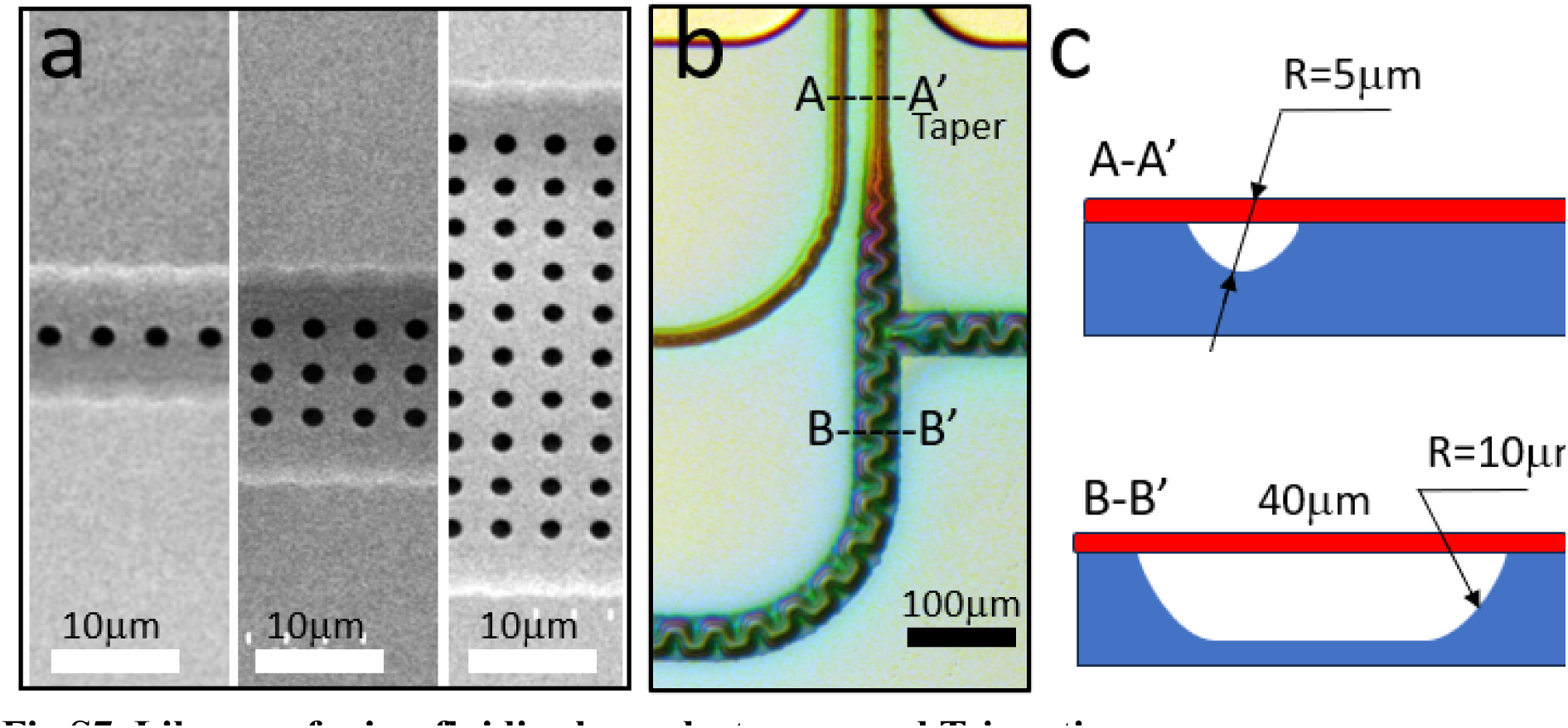
Library of microfluidic channels, tapers, and T-junctions. A) SEM microphotograph of 3 microfluidic channels of different width defined lithographically by number of rows of perforations. B) Optical microphotograph of an area of T-junction with narrow (top) and wide (bottom) channels connected with a gradual taper. C) Schematics of cross-sections of the narrow (top) and wide (bottom) channels.

